# Selectively expressing SARS-CoV-2 Spike protein S1 subunit in cardiomyocytes induces cardiac hypertrophy in mice

**DOI:** 10.1101/2021.06.20.448993

**Authors:** Steven G. Negron, Chase W. Kessinger, Bing Xu, William T. Pu, Zhiqiang Lin

## Abstract

Cardiac injury is common in hospitalized COVID-19 patients and portends poorer prognosis and higher mortality. To better understand how SARS-CoV-2 (CoV-2) damages the heart, it is critical to elucidate the biology of CoV-2 encoded proteins, each of which may play multiple pathological roles. For example, CoV-2 Spike glycoprotein (CoV-2-S) not only engages ACE2 to mediate virus infection, but also directly impairs endothelial function and can trigger innate immune responses in cultured murine macrophages. Here we tested the hypothesis that CoV-2-S damages the heart by activating cardiomyocyte (CM) innate immune responses. HCoV-NL63 is another human coronavirus with a Spike protein (NL63-S) that also engages ACE2 for virus entry but is known to only cause moderate respiratory symptoms. We found that CoV-2-S and not NL63-S interacted with Toll-like receptor 4 (TLR4), a crucial pattern recognition receptor that responsible for detecting pathogen and initiating innate immune responses. Our data show that the S1 subunit of CoV-2-S (CoV-2-S1) interacts with the extracellular leucine rich repeats-containing domain of TLR4 and activates NF-kB. To investigate the possible pathological role of CoV-2-S1 in the heart, we generated a construct that expresses membrane-localized CoV-2-S1 (S1-TM). AAV9-mediated, selective expression of the S1-TM in CMs caused heart dysfunction, induced hypertrophic remodeling, and elicited cardiac inflammation. Since CoV-2-S does not interact with murine ACE2, our study presents a novel ACE2-independent pathological role of CoV-2-S, and suggests that the circulating CoV-2-S1 is a TLR4-recognizable alarmin that may harm the CMs by triggering their innate immune responses.

## Introduction

The COVID-19 pandemic has resulted in more than 170 million COVID-19 cases and 3.5 million deaths. Before its ending and following its long-term consequences, additional toll has yet to be tallied. SARS-CoV-2 (CoV-2) infects not only the respiratory tract but also the other major organs, including the heart (Puelles et al., 2020). In hospitalized COVID-19 patients, cardiac injury portends higher mortality rate (Guo et al., 2020; Mitrani et al., 2020) and poorer prognosis (Fayol et al., 2021). In recovered COVID-19 patients who experienced myocardial injury, cardiac function abnormalities have been observed (Fayol et al., 2021; Wang et al., 2021), raising the concern as to whether CoV-2 infection causes permanent cardiac damages and leads to chronic cardiac complications (Yancy and Fonarow, 2020; Shchendrygina et al., 2021).

Although the underlying mechanisms of COVID-19 cardiac injury are obscure, several hypotheses have been proposed: CoV-2 infection of CMs leads to myocarditis(Bulfamante et al., 2020; Dolhnikoff et al., 2020); CoV-2 infection of endothelial cells results in vascular leakage and microvascular thrombi; exaggerated systemic inflammation causes cytokine storm which stresses the heart (Akhmerov and Marban, 2020); and severe pneumonia can induce multiple organ failure(Jirak et al., 2021). Ultimately, all of these different mechanisms converge to damage the main functional units of the heart: the cardiomyocytes (CMs).

Similar to professional innate immune leukocytes, CMs have their own innate immune machinery (Brown, 2020), activation of which is beneficial for defending CMs against pathogens invasion but may cause cardiac inflammation and myocardial damage (Hobai et al., 2015; Brown, 2020). One of the best-characterized innate immune pathways is the Toll-like receptor 4 (TLR4)/NF-kB axis, which also plays essential roles in cardiac inflammation and hypertrophic remodeling (Yang et al., 2016). Under pathologic conditions, such as septic shock or myocardial infarction, depletion of TLR4 in CMs attenuates cardiac injury (Fallach et al., 2010). On the contrary, activation of TLR4 in cultured CMs impairs their contractility and induces the expression of pro-inflammatory cytokine genes (Boyd et al., 2006).

To better understand how CoV-2 damages the heart, it is critical to elucidate the biology of CoV-2 encoded proteins, each of which may play multiple pathological roles. For example, CoV-2 Spike glycoprotein (CoV-2-S) not only engages angiotensin-converting enzyme 2 (ACE2) to mediate virus infection (Zhou et al., 2020), but also directly impairs endothelial function (Lei et al., 2021) and triggers TLR4/NF-kB signaling in cultured macrophages (Shirato and Kizaki, 2021; Zhao et al., 2021). ACE2 is a crucial enzyme of the renin angiotensin system (RAS) that maintains blood pressure homeostasis (Teerlink, 1996), and is critical in modulating cardiac hypertrophy through Angiotensin 2 regulation. In COVID-19 patients, CoV-2-S may impair cardiac ACE2 function and expression, as it does in endothelial cells (Lei et al., 2021), therefore leading to cardiac injury (Chen et al., 2020a). Alternatively, it may also damage the heart through ACE2-independent mechanisms.

Here we investigated the contribution of CoV-2-S activation of TLR4/NK-κB to cardiac inflammation and dysfunction. We found that CoV-2-S directly interacted with TLR4 and activated NF-kB transcriptional activity. We also found that selectively expressing truncated CoV-2-S in CMs caused heart dysfunction, inflammation and hypertrophic remodeling. Since CoV-2-S does not interact with murine ACE2 (Zhou et al., 2020), our study presents a novel ACE2-independent pathological role CoV-2-S.

## Results

### CoV-2-S interacts with TLR4 to activate NF-kB

Until now, seven human coronaviruses (HCoVs) have been identified (Figure 1A), including 2 αCoV (229E and NL63) and 5 βCoV (OC43 and HKU1 [lineage A], SARS-CoV and SARS-CoV-2 [lineage B], MERS-CoV [lineage C]). Among them, four (229E, NL63, OC43 and HKU1) cause minor to moderate respiratory tract infections, and three (MERS, SARS-CoV, SARS-CoV-2) cause severe acute respiratory syndrome (Chan et al., 2020).

**Figure 1.**
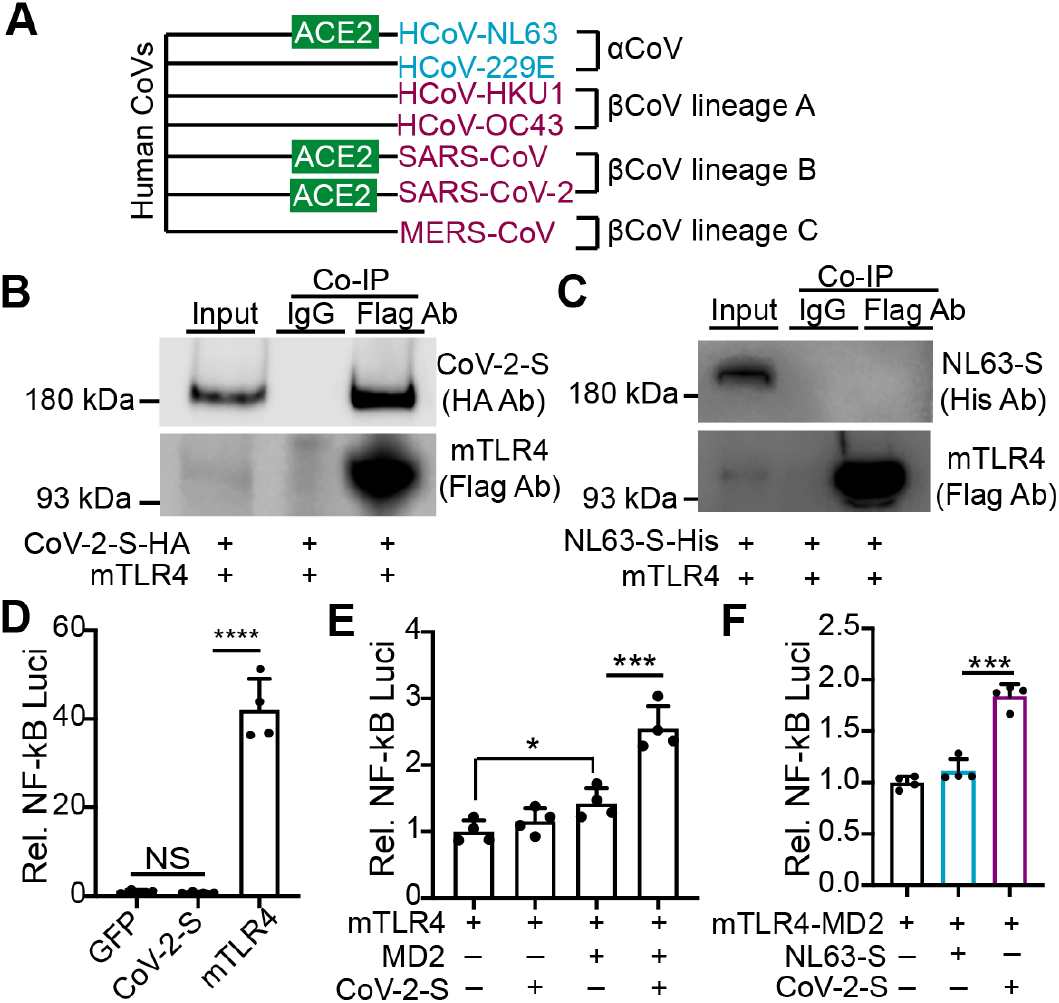
SARS-CoV-2 Spike protein interacts with TLR4 to activate NF-kB. **A**. Taxonomy of human Coronavirus (CoV). HCoV-NL63, SARS-CoV, and SARS-CoV-2 Spike proteins use ACE2 to mediate virus entry. **B-C**. Co-immunoprecipitation (Co-IP) assays. Plasmids expressing FLAG tagged murine TLR4 (mTLR4) was transfected into HEK293T cells together with plasmids expressing HA tagged SARS-CoV-2 Spike (CoV-2-S, B) and 8xHis tagged HCoV-NL63 Spike (NL63-S, C), respectively. One day after transfection, cells were collected for FLAG antibody Co-IP. Mouse IgG was used as negative control. **D-F**. NF-kB luciferase report assay. Luciferase reporter containing 4xNF-kB binding sites (NF-kB Luci) was co-transfected with indicated plasmids into HEK293T cells. One day after transfection, cells were collected for luciferase activity test. One-Way ANOVA test, *, P<0.05, ***, P<0.001; **** P<0.0001. N=4.

Although both NL63 and SARS-CoV-2 (CoV-2) use ACE2 as their receptor to infect host cells, NL63 is much less virulent than CoV-2, indicating that tissue tropism may not account for their dramatic virulent differences. Because CoV-2 but not NL63 causes severe cytokine storm (Mehta et al., 2020), and two independent studies have shown that CoV-2 Spike protein (CoV-2-S) activates TLR4/NF-kB signaling in macrophages (Zhao et al., 2021), we then suspected that CoV-2-S and NL63-S might have different effects on TLR4/NF-kB signaling.

To address this issue, we first performed co-immunoprecipitation (Co-IP) assays. In HEK293T cells, Flag tagged murine TLR4 (mTLR4) successfully pulled down CoV-2-S and did not pull down NL63-S (Fig. 1B-C), confirming that CoV-2-S and not NL63-S interacts with TLR4. Second, we used a 4xNF-kB Luciferase reporter (NF-kB_Luci) to determine whether CoV-2-S and NL63-S differentially regulates TLR4/NF-kB signaling axis in HEK293T cells. To initiate downstream signaling, TLR4 needs to form a heterozygous complex with MD2 (Dejkhamron et al., 2007), and these two proteins are both absent in HEK293T cells (Medvedev and Vogel, 2003). In line with the published data, NF-kB_Luci was active only when it was co-expressed with TLR4 (Fig. 1D), and TLR4 activation of NF-kB was further enhanced by adding in MD-2 (Fig. 1E). In the absence of TLR4 and MD-2, CoV-2-S did not affect NF-kB reporter activity (Fig. 1D). CoV-2-S did not stimulate NF-kB reporter activity in the presence of TLR4; however, it enhanced NF-kB transcriptional activity when it was co-expressed with TLR4 and MD-2 (Fig. 1E). Different from CoV-2-S, NL63-S did not affect NF-kB transcriptional activity in the presence of TLR4 and MD2 (Fig. 1F).

These data demonstrate that CoV-2-S directly interacts with TLR4, and suggest that CoV-2-S activation of NF-kB depends on TLR4/MD-2 complex. Given that CoV-2 and not HCoV-NL63 causes cytokine storm, and that CoV-2-S but not NL63-S activates TLR4/NF-kB signaling, the current data suggest that CoV-2-S activation of TLR4/NF-kB signaling may be one of the underlying mechanisms that account for the high virulence of CoV-2.

### The S1 subunit of CoV-2 Spike protein (CoV-2-S1) interacts with the extracellular Leucine-rich repeat region of TLR4

CoV-2-S has two subunits: S1 and S2. S1 contains receptor binding domain (RBD), which binds ACE2 (Fig. 2A). S2 contains functional domains responsible for virus and host cell membrane fusion (Shang et al., 2020). S1 and S2 are connected by a protein convertase site (PPC), which can be cleaved by host proteases (Walls et al., 2020). To further characterize the relationship between CoV-2-S and TLR4, we generated different constructs to map the interacting regions of these two proteins. We first generated constructs expressing different CoV-2-S subunits (Fig. 2A). Co-IP results showed that CoV-2-S1 and not CoV-2-S2 was pulled down by TLR4 (Fig. 2B-C). TLR4 has three essential domains: extracellular Leucin rich repeat (LRR) domain (Fig. 2D), transmembrane domain (TM) and intracellular Toll/In-terleukin-1 receptor like domain (TIR) (Rock et al., 1998; Vaure and Liu, 2014). We then generated constructs expressing either TLR4-LRR or TLR4-TIR (Fig. 2D). Co-IP results showed that TLR4-LRR but not TLR4-TIR pulled down CoV-2-S1 (Fig. 2E). Together with Figure 1, these data demonstrate that CoV-2-S1 directly interacts with the LRR domain of TLR4. In COVID-19 patients, CoV-2-S1 can be shed into blood stream, and the serum levels of CoV-2-S1 are associated with COVID-19 severity (Ogata et al., 2020). These data suggest that circulating CoV-2-S1 may serve as an alarmin to activate host innate immune responses through TLR4 (Fig. 2F).

**Figure 2.**
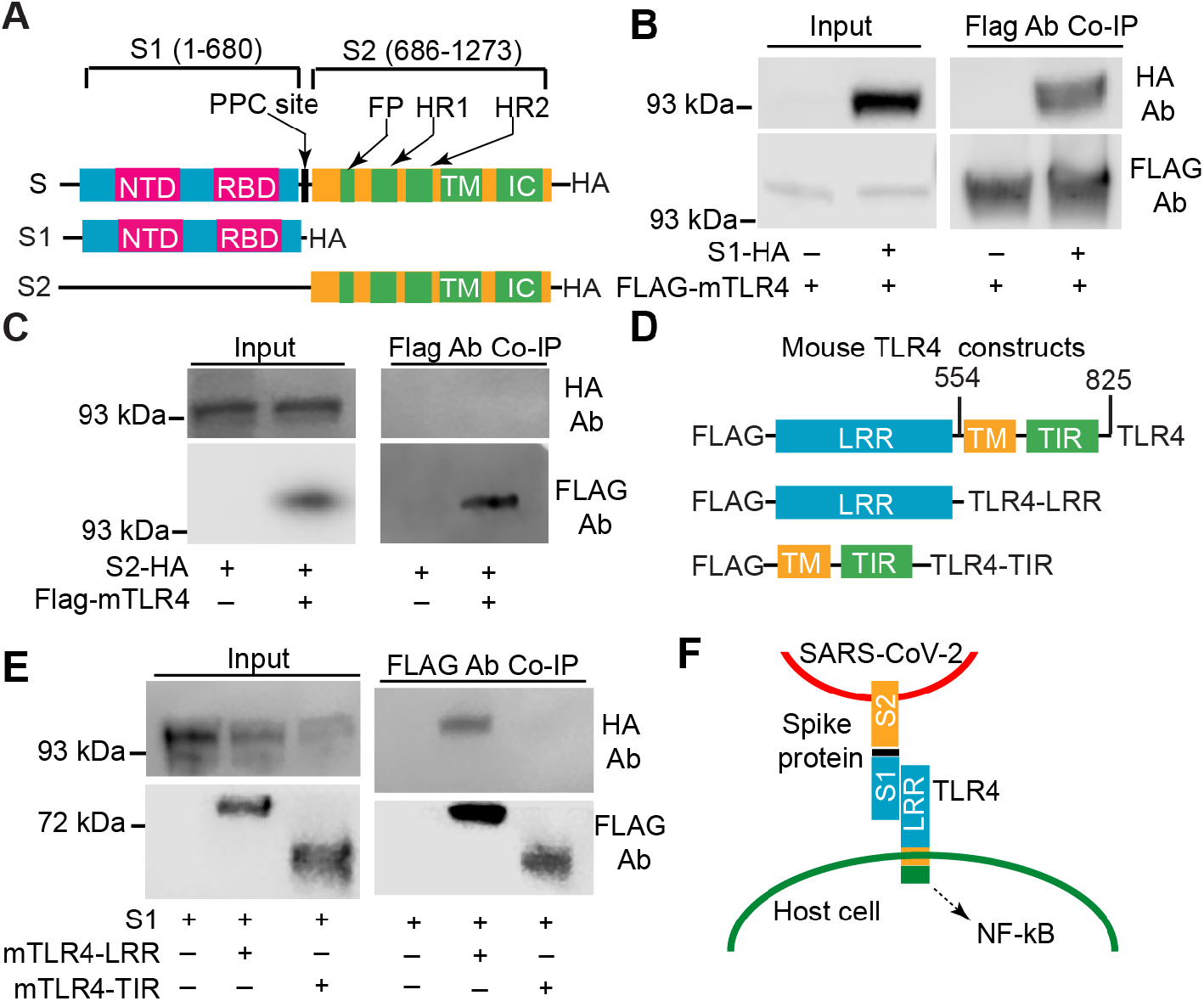
SARS-COV-2 Spike protein S1 subunit interacts with the Leucine-rich repeat region of TLR4. **A**. Schematic view of constructs expressing SARS-CoV-2 Spike protein and its different subunits. S1, receptor binding subunit; S2, membrane fusion subunit. NTD, N-terminal domain. RBD, receptor binding domain. PPC, proprotein convertase (PPC) motif. FP, fusion peptide. HR1, heptad repeat 1. HR2, heptad repeat 2. TM, transmembrane anchor. IC, intracellular tail. HA, hemagglutinin epitope tag. FP, HR1 and HR2 domains mediate membrane fusion between SARS-CoV-2 and host cells. **B-C**. SARS-CoV-2 Spike protein interacts with TLR4 through its S1 subunit. **D**. Constructs expressing TLR4 and its different functional domains. LRR, Leucine-rich repeat region. TM, transmembrane domain. TIR, Toll/Interleukin-1 receptor domain. **E**. SARS-CoV-2 Spike protein S1 subunit interacts with the LRR domain of TLR4. B,C,E, indicated plasmids were transfected into HEK293T cells. One day after transfection, cells were collected for FLAG antibody Co-IP. **F**. Diagram summary of the relationship between SARS-CoV-2 Spike protein and host cell TLR4.

### Membrane-localized CoV-2-S1 is sufficient to activate TLR4/NF-kB signaling

Since the outbreak of COVID-19 pandemic, several experimental systems have been developed to study the pathophysiology of CoV-2 in CMs. *In vitro*, CoV-2 has been used to infect human induced pluripotent stem cell-derived cardiomyocytes (iPS-CMs) or cardiac organoids (Mills et al., 2021). CoV-2 infection of iPS-CMs triggered innate immune responses and induced apoptosis (Bojkova et al., 2020; Chen et al., 2020b; Sharma et al., 2020); however, it is unknown how CoV-2 damages CMs. One potential mechanism is that CoV-2-S injures CMs by triggering their innate immune responses. In order to test this hypothesis *in vivo*, we determined to use an AAV9 vector to express CoV-2-S in the mouse heart. To reach this goal, three hurdles needed to be surmounted. First, at 3.8 kb, the CoV-2-S coding sequence (Ogawa et al., 2020) is too large to be efficiently expressed by AAV9, therefore a coding sequence that expresses a truncated yet functional CoV-2-S needs to be generated. Second, CoV-2-S can be cleaved at the PPC site, thus shedding S1 subunit into extracellular space (Zhang et al., 2020), and this S1 shedding potentially impedes the efforts of deciphering CoV-2-S cell-autonomous effects. Third, to mimic the sub-cellular localization of the full length CoV-2-S(Bojkova et al., 2020), a transmembrane domain needs to be included in the truncated CoV-2-S.

To overcome these barriers, we first generated a construct to express a truncated, membrane localized CoV-2-S (pCAG.S1-TM, Fig. 3A). This engineered S1-TM retains the S1 subunit, the transmembrane anchor (TM) and intracellular tail (IC) domains, and does not contain the PPC site, FP and HR domains (Fig. 3A). Co-IP confirmed that S1-TM interacted with TLR4 (Fig. 3B), and NF-kB luciferase reporter assay indicated that S1-TM activation of NF-kB was not distinguishable from that of CoV-2-S1 or full length CoV-2-S (Fig. 3C). Additionally, in HEK293T cells, we confirmed that a portion of S1-TM was localized on plasma and nuclear membranes (Fig. 3D). These data indicate S1-TM can be used for dissecting CoV-2-S innate immune effects.

**Figure 3.**
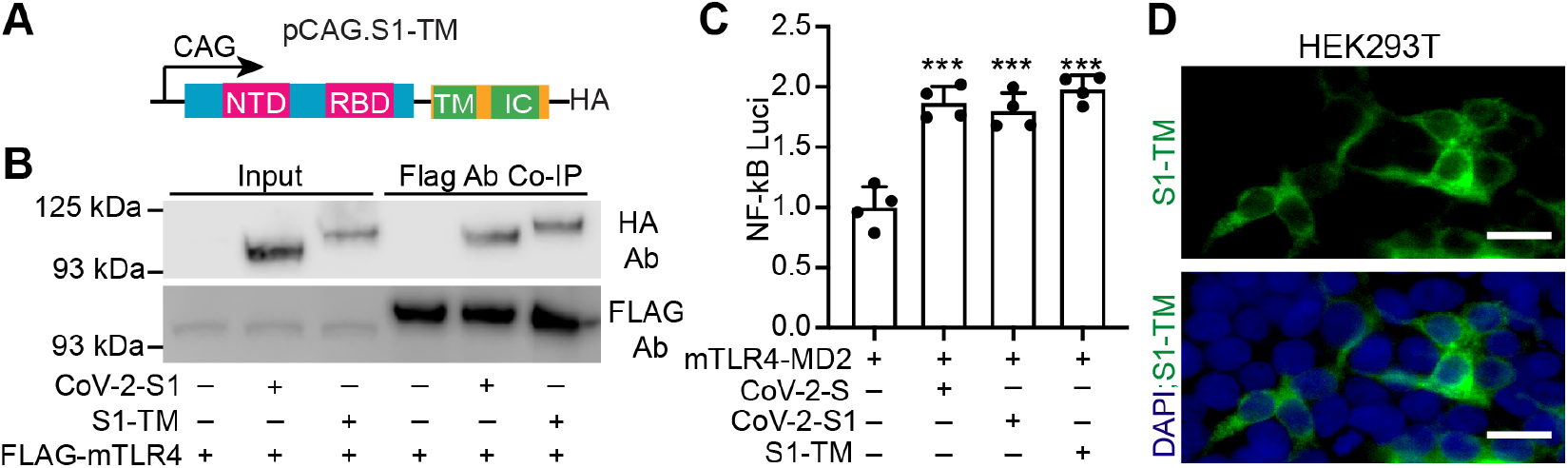
Truncated CoV-2 Spike protein containing S1 subunit and transmembrane anchor domain is sufficient to activate NF-kB. **A**. Schematic view of a construct expressing truncated SARS-CoV-2 Spike protein. S1 subunit, transmembrane anchor domain and intracellular tail were cloned in frame into pCAG-HA plasmid. The expression of the truncated Spike protein (S1-TM) was driven by the constitutively active CAG promoter. **B**. Co-IP assay. Indicated plasmids were transfected into HEK293T cells. One day after transfection, cells were collected for Co-IP. **C**. NF-kB luciferase report assay. NF-kB Luci plasmid was co-transfected with indicated plasmids into HEK-293T cells. One day after transfection, cells were collected for luciferase activity test. One-Way ANOVA test, ***, P<0.01. N=4. **D**. Sub-cellular localization of S1-TM. Scale bar = 50 μm.

### Expressing S1-TM in the heart reduces cardiac systolic function

To study the innate immune effects of CoV-2-S in mouse hearts, we cloned S1-TM into our cardiac specific AAV vector (Lin et al., 2014) and generated AAV9.cTnT.S1-TM (Fig. 4A). As illustrated in Fig. 4B, AAV9.cTnT.S1-TM (AAV.S1-TM), or AAV9.cTnT.GFP (AAV.GFP) control, was delivered retro-orbitally at a dose of 4 × 10^10^ vg/g to three-week-old mice. Two weeks after AAV delivery, heart function was evaluated via M-mode echocardiography, after which mice were sacrificed for heart collection.

**Figure 4.**
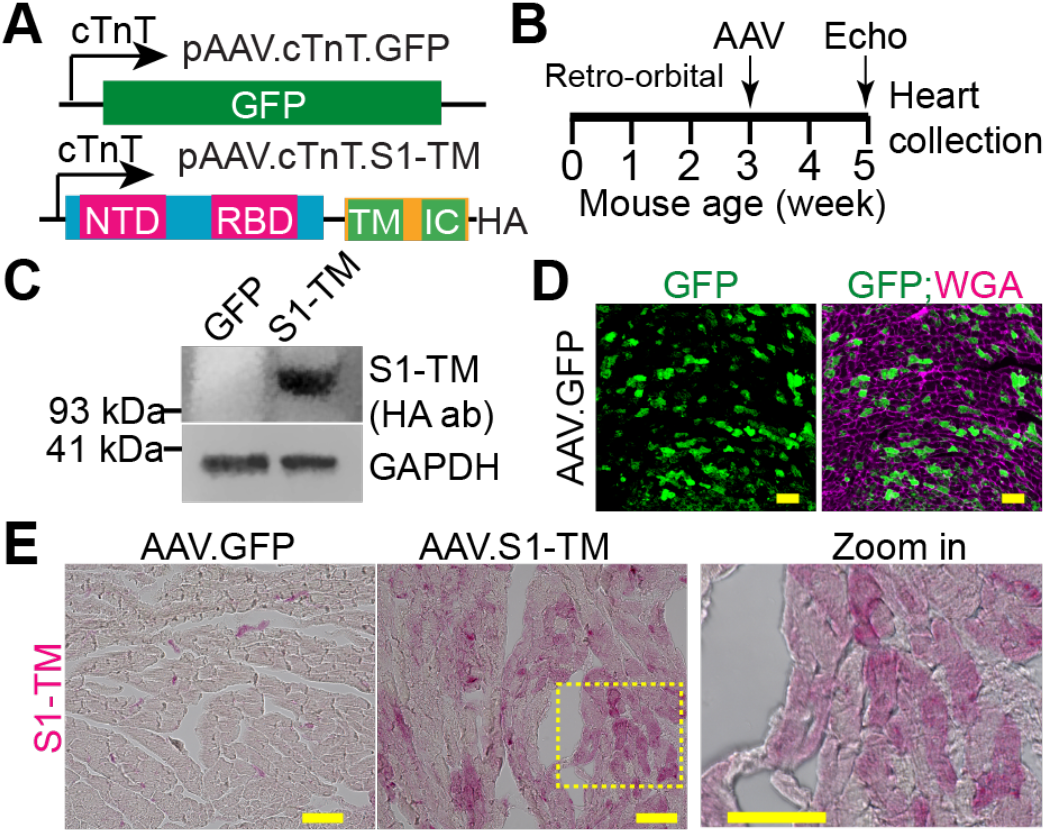
Generation and validation of AAV9.cTnT.S1-TM. **A.** Cardiac specific AAV constructs. The expression of cargo genes is driven by chicken cardiac Troponin T (cTnT) promoter. **B**. Experimental design. **C**. Immuno Blot of S1-TM. **D**. GFP and wheat germ agglutinin (WGA) stained myocardium. **E**. Immunohistochemistry images of Spike protein antibody stained myocardium. Alkaline phosphatase based detection system was used to visualize the expression of S1-TM. D, E, Scale bar = 50 μm.

Western blot confirmed that S1-TM was expressed in the AAV.S1-TM hearts (Fig. 4C). As reflected by GFP and S1-TM staining, the dose of 4 × 10^10^ vg/g achieved ~30-40% CM trans-duction (Fig. 4D, 4E). Furthermore, immunohistochemistry staining demonstrated that S1-TM was expressed in CMs (Fig. 4E). These data confirm that AAV.S1-TM efficiently transduces the myocardium and specifically expresses S1-TM in CMs.

During the two-week monitoring time window, AAV.S1-TM-transduced mice had no noticeable weight loss and behaved normally. As mentioned above, M-mode echocardiography was performed with conscious mice two weeks after AAV injection. Compared to AAV.GFP control mice, both male and female AAV.S1-TM mice had moderately decreased fractional shortening values (Fig. 5A, 5B). The heart rate of these mice were similar (Fig. 5C), indicating that S1-TM dose not affect heart rhythm. These data suggest that expressing S1-TM in CMs decreases systolic heart function.

**Figure 5.**
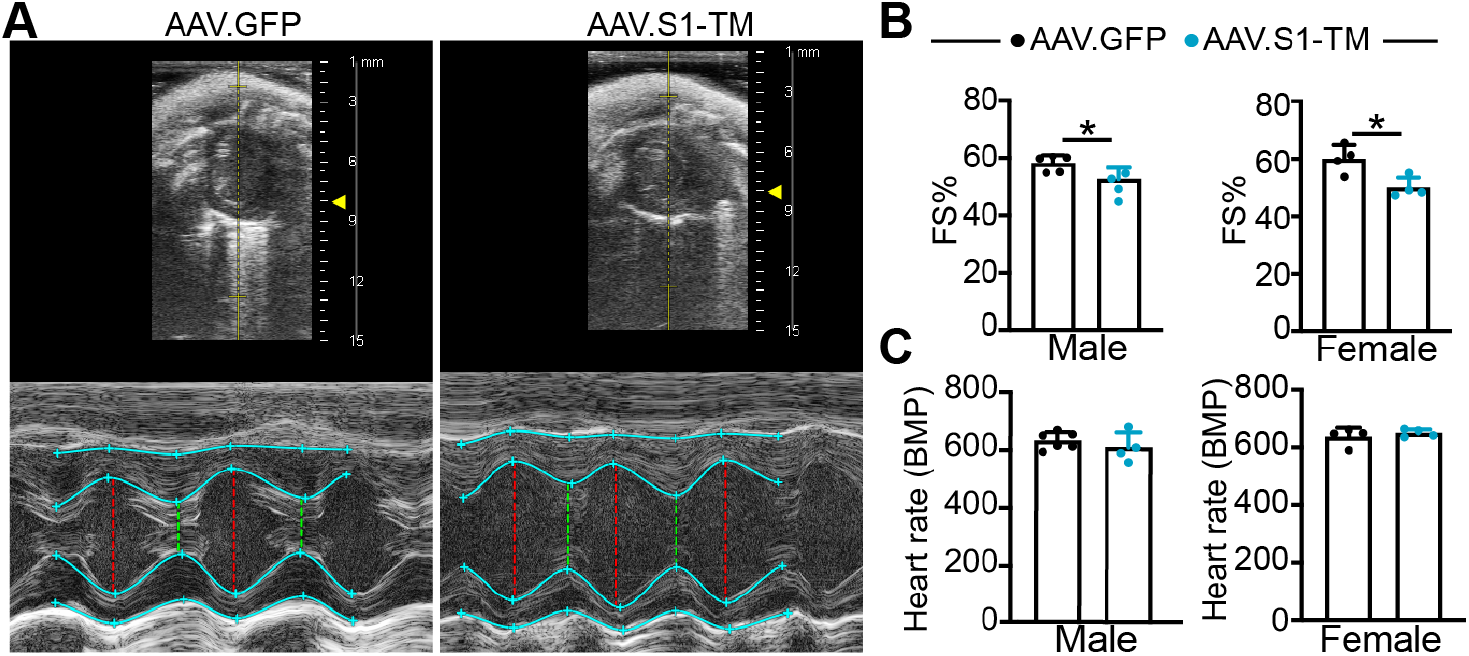
S1-TM reduces heart function. **A.** M-mode echocardiograms at papillary muscle level. Top panel, short axis view of the left ventricle; bottom panel, M-mode tracing of the left ventricle. Echocardiography was performed with conscious mice. **B-C**. Echocardiography measurement of heart systolic function (B) and heart rate (C). B, Student’s t-test, *, P<0.05. B-C, n= 4-5.

### Expressing S1-TM in the heart induces cardiac hypertrophy

Two weeks after AAV delivery, the collected hearts were examined for morphological and molecular differences. AAV.S1-TM mice had a higher heart-to-body weight ratio (Fig. 6A) and larger hearts (Fig. 6B-C) than AAV.GFP mice. Furthermore, CMs were enlarged in AAV.S1-TM hearts (Fig. 6D-E). In AAV.S1-TM hearts, *Myh6* was substantially reduced. Additionally, *Nppa*, a cardiac hypertrophy marker gene, was robustly up-regulated by S1-TM. These data suggest that S1-TM induces cardiac stress responses that result in cardiac hypertrophic remodeling.

**Figure 6.**
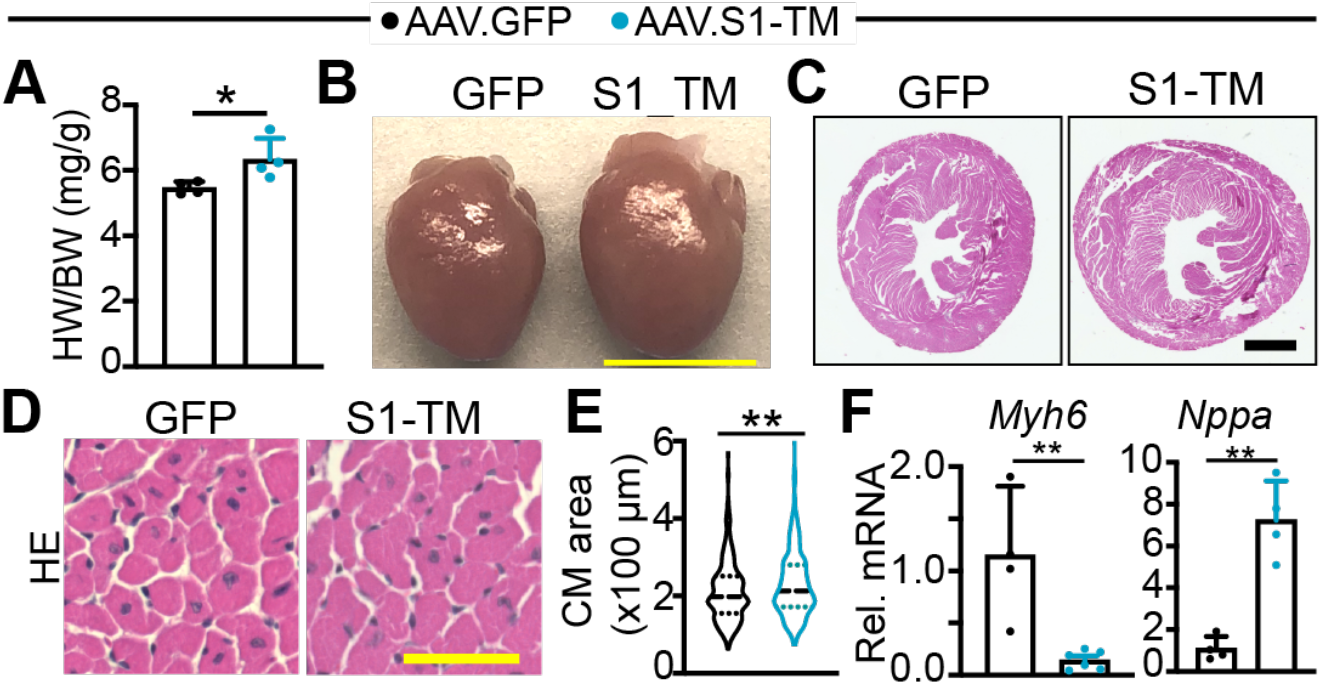
S1-TM causes cardiac hypertrophy. **A**. Heart-to-body weight ratio. N=4. Student’s t test, *, P<0.05. **B**. Heart gross morphology. Scale bar = 5 mm. **C**. Heart cross section stained by Hematoxylin and Eosin (HE). Scale bar = 1mm. **D**. Myocardium stained by HE. Scale bar = 50 μm. **E**. CM cross-section area measurement. In each group, CMs from three different hearts were measured. AAV.GFP, n= 358. AAV.S1-TM, n=396. Mann-Whitney test, **, P<0.01. F. qRT-PCR measurement of Myh6 and Nppa. N=4-6. Student’s t test, **, P<0.01.

### Expressing S1-TM in the heart induces cardiac inflammation

Histological analysis of AAV.S1-TM myocardium stained with hematoxilyn and eosin (H&E) showed regions enriched with small and round-shape cells (Fig. 7A), suggesting that expressing S1-TM in CMs may induce cardiac inflammation. Infiltration of activated macrophages is associated with inflammation, and Mac-3 is a marker for activated macrophages (Frantz and Nahrendorf, 2014). Mac-3 immunohistochemistry staining of the AAV.S1-TM myocardium revealed that activated macrophages were populating in the lesion regions (Fig. 7B). H&E staining of AAV.GFP myocardium showed intact myocardium (Fig. 7A), and Mac-3 staining of these hearts revealed no to minimal presence of activated macrophages (Fig. 7B).

**Figure 7.**
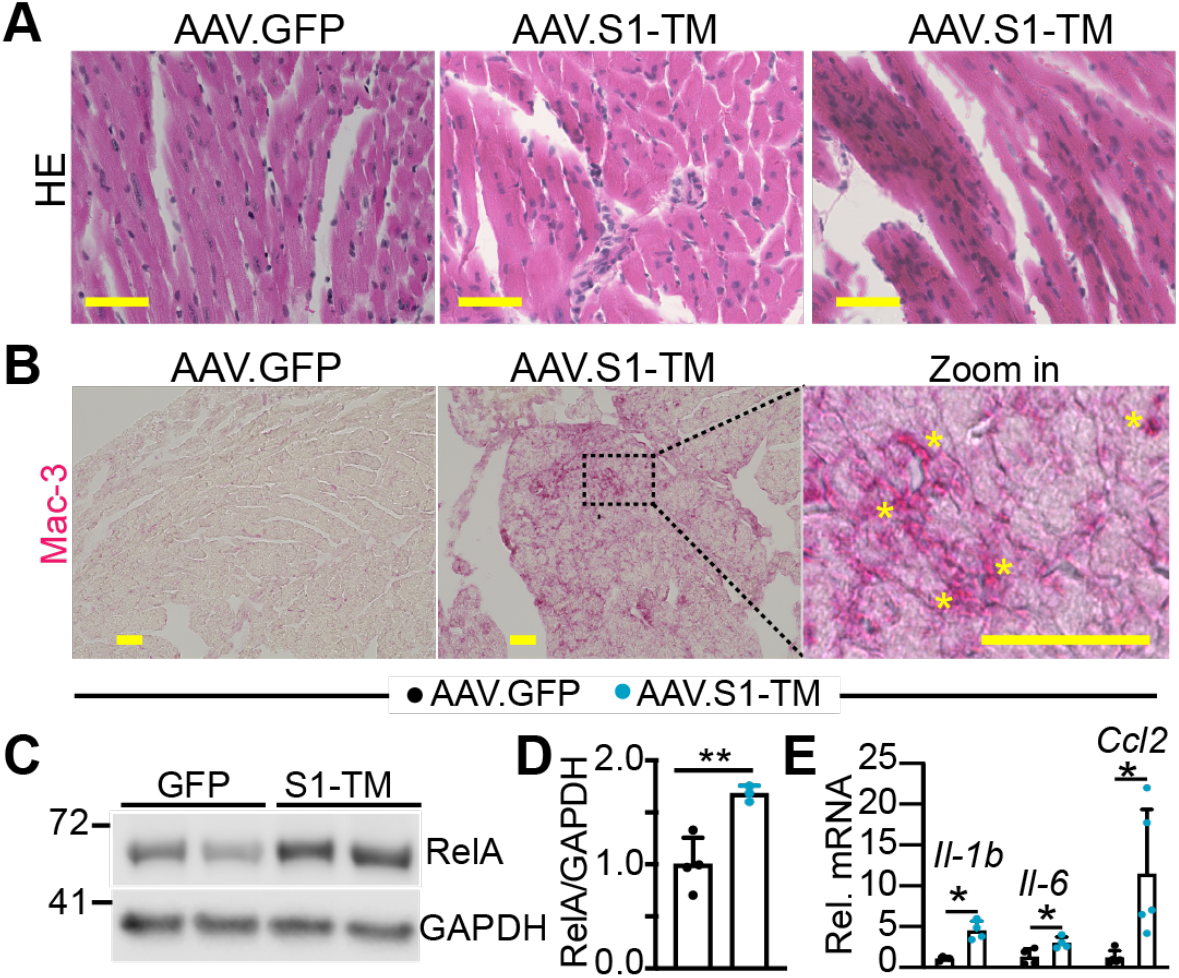
S1-TM induces cardiac inflammation. **A**. HE stained myocardium. Scale bar = 50μm. **B**. Immunohistochemistry staining of Mac-3 in myocardium. Alkaline phosphatase based detection system was used to visualize Mac-3 positive cells. In the zoom-in image, yellow stars indicate macrophage clusters. Scale bar = 50μm. **C**. Immuno Blot of RelA. **D**. Densitometry quantification of RelA. RelA protein levels were normalized to GAPDH. N=3-4. **E**. qRT-PCR measurement of Il-1b, Il-6 and Ccl2. N=4-5. D,E, Student’s t test, *, P<0.05; **, P<0.01.

These histological data strongly suggest that expressing S1-TM in the CMs induces cardiac inflammation. To further confirm this hypothesis on molecular level, we examined RelA protein expression and measured the mRNAs of two pro-inflammatory cytokine genes*: Il-1b* and *Il-6*. Compared to AAV.GFP hearts, AAV.S1-TM hearts had significantly higher expression of RelA, *Il-1b* and *Il-6* (Fig. 7C-E). *Ccl2*, a crucial chemokine gene whose protein product facilitates innate immune cells infiltration (Dewald et al., 2005), was also robustly up-regulated in the AAV.S1-TM hearts (Fig. 7E). Altogether, these data suggest that expressing S1-TM in the CMs triggers cardiac inflammation.

## Discussion

### CoV-2 Spike protein (CoV-2-S) is recognized by TLR4

In critically ill COVID-19 patients, besides acute respiratory distress, systemic inflammation is another major clinical manifestation. Based on *in silico* analysis, a superantigen hypothesis has been proposed to explain how CoV-2 infection may cause systemic inflammation: *CoV-2-S* contains a superantigen motif near its S1/S2 cleaveage site. In toxic shock syndrome, this motif interacts with both T cell receptor (TCR) and CD28 to elicit T lymphocyte activation (Cheng et al., 2020). Two independent *in vitro* studies have shown CoV-2-S to activate TLR4 innate immune signaling in cultured macrophages (Shirato and Kizaki, 2021; Zhao et al., 2021). These studies highlight CoV-2-S as a potent culprit that could possibly induce systemic inflammation; however, the underlying molecular mechanism is not fully understood. CoV-2-S has two subunits, with its S1 subunit (CoV-2-S1) being able to be shed into blood stream (Ogata et al., 2021), Importantly, in COVID-19 cases, the serum levels of CoV-2-S1 shows higher positive correlation with clinical severity than CoV-2 nucleocapsid protein (Ogata et al., 2020). Here, our data show that CoV-2-S1 interacts with the extracellular LRR domain of TLR4 and activates NF-kB, suggesting that soluble CoV-2-S1 shed by CoV-2 or infected host cells is a TLR4-recognizable alarmin. In summary, CoV-2 infection may cause systemic inflammation through three different ways: i) virus infection of host cells leads to cell death and activation of innate immune responses; ii) the spike protein complex protruding out of the virus envelope serves as a superantigen to elicit T cell-mediated inflammation; iii) circulating CoV-2-S1 exacerbates inflammation through its activation of innate immune responses.

### CoV-2-S is toxic to CMs

CoV-2 can damage the heart by directly infecting the CMs and therefore causing myocarditis (Bulfamante et al., 2020; Dolhnikoff et al., 2020), it can also injure the heart indirectly (Nishiga et al., 2020). In either senario, CMs are potentially exposed to CoV-2-S originating from different sources, such as virus particles and CoV-2-S1 shed by virus or infected cells (Zhang et al., 2020).

Currently, small animal models of COVID-19, including hamster and K8-hACE2 transgenic mice (Bao et al., 2020; Chen et al., 2020b), cause severe weight loss (Mills et al., 2021) and may potentially damage multiple organs, precluding a clear understanding of the specific cellular toxicity and/or innate immune responses related to CoV-2-S/S1. To define the pathological role of CoV-2-S in mouse hearts, we chose to use the AAV9 vector to selectively express truncated CoV-2-S (S1-TM) in CMs. This engineered S1-TM retains the S1 subunit, the transmembrane anchor (TM) and intracellular tail (IC) domains, but does not contain the PPC site or the membrane fusion functional domains, thus allowing us to pinpoint the cell-autonomous effects of CoV-2-S. Our data demonstrate that selectively expressing S1-TM in CMs injures the heart and induces cardiac inflammation, suggesting that CoV-2-S1 is toxic to the CMs. Given that CoV-2-S1 interacts with TLR4 and activates NF-kB transcriptional activity *in vitro*, and expressing S1-TM in the heart activates NF-kB target genes, it is likely that CoV-2-S1 harms the CMs by activating their innate immune responses.

### CoV-2-S harms CMs independent of ACE2 in mouse hearts

The renin angiotensin system (RAS) plays central roles in maintaining the blood pressure homeostasis (Teerlink, 1996). In this system, renin cleaves Angiotensiongen to produce Angiotensin (inactive), which is further converted to pro-inflammatory and vasoconstrictive Angiotensin II (Ang II) by Angiotensin converting enzyme (ACE). As a homologue of ACE, ACE2 antagonizes ACE function by converting AngII to Ang 1-7 (anti-inflammatory and vasodilatory). Since CoV-2-S engages ACE2 to mediate virus entry, it may harm CMs by destabilizing ACE2 (Chen et al., 2020a), as it does in endothelial cells (Lei et al., 2021). Nevertheless, emerging evidence suggests that CoV-2-S may also harm the heart through ACE2-independent mechanisms. First, both HCoV-NL63 and SARS-CoV-1/2 use human ACE2 (Li et al., 2007) for cellular entry, but only SARS-CoV-1/2 infection causes cardiac injury. Second, CoV-2-S interacts with cellular heparan sulfate (Clausen et al., 2020) and may activate the alternative complement pathway (Yu et al., 2020). Third, CoV-2-S activates inflammatory responses in murine macrophages (Shirato and Kizaki, 2021), which express CoV-2-S incompatible ACE2 (Zhou et al., 2020). Here, we provided direct evidence suggesting the existence of an ACE2-independent pathogenesis mechanism of COVID-19 associated cardiac injury: CoV-2-S but not NL63 Spike protein interacts with TLR4 and activates NF-kB (Fig. 1); selectively expressing truncated CoV-2-S in CMs injures the mouse hearts and activates the expression of NF-kB target genes. We are now underway in testing whether blunting TLR4/NF-kB signaling protects the heart against CoV-2-S stress.

### Limitation of the current study

In severe COVID-19 patients, cardiac injury is likely a result of acute pneumonias (Jirak et al., 2021; Metkus et al., 2021). Since our AAV only express S1-TM in the CMs, our system is not suitable for studying the pathogenesis of severe COVID-19 related cardiac injury. Rather, this AAV system can be used for investigating the pathogenesis of myocardial injury without COVID-19 acute respiratory distress, and it is also a powerful platform for dissecting the pathological roles of CoV-2 encoded genes in the heart or other organs. In addition, the current study did not test whether CoV-2-S impairs CMs electrophysiology, a crucial question to be addressed in future studies.

## Materials and methods

Supplemental information provides expanded material and experimental procedures.

### Experimental animals

All animal procedures were approved by the Institute Animal Care and Use Committees of Masonic Medical Research Institute. All experiments were performed in accordance with NIH guidelines and regulations.

### Luciferase reporter assay

HEK 293T cells were cultured in 24 well plates for luciferase assay. 100 ng/well indicated plasmids and 10 ng pRLTK internal control vector (Promega) were co-transfected with 1.25 μl Lipofectamine 3000 (Invitrogen). Luciferase activity was measured 24 hours after transfection using the Dual-Luciferase reporter assay system (Promega).

### Gene Expression

Total RNA was isolated using Trizol. For qRT-PCR, RNA was reverse transcribed (Applied Biological Materials Inc., G454) and specific transcripts were measured using Sybr Green chemistry (Lifesct., LS01131905Y) and normalized to *Gapdh*. Primer sequences were provided in Supplementary Table 2. Primary antibodies used for immuno blot and immunohistochemistry staining were listed in Supplementary Table 3.

### Statistics

Values were expressed as mean ± SD. Student’s t-test or ANOVA with Tukey’s honestly significant difference post hoc test was used to test for statistical significance involving two or more than two groups, respectively.

## Sources of Funding

Z.L. was supported by the Masonic Medical Research Institute.

## Conflict of interest

none declared

## Notes

### Competing Interest Statement

The authors have declared no competing interest.

