## Supplemental Information for "Selectively expressing SARS-CoV-2 Spike protein S1 subunit in cardiomyocytes induces cardiac hypertrophy in mice"

### Material and Methods

#### Plasmids

SARS-CoV-2 Spike-HA (#141347), HCoV-NL63 Spike-8xHis (#166017), mTLR4 (#13087), hMD2 (#13028), NF-kB luciferase reporter (#111216) plasmids were all purchased from Addgene. AAV plasmids: The coding sequences of SARS-CoV-2 Spike S1 and S2-HA were PCR amplified from plasmid #141347, and were cloned into pAAV.cTnT.vector (Lin et al., 2014), respectively. Annealed DNA oligos coding HA tag was cloned in-frame into pAAV.cTnT.S1 with NcoI and SalI, yielding out pAAV.cTnT.S1-HA. To generate pAAV.cTnT.S1-TM, the transmembrane and intracellular tail domains of S2 was amplified by PCR, and cloned in-frame to pAAV.cTnT.S1 with NcoI and SalI. To have a high expression efficiency, the different Spike protein coding sequences were further cloned into pAAV.CAG vector (Mao et al., 2011) following two cloning steps: i) pAAV.cTnT.S1-HA, pAAV.cTnT. S2-HA, and pAAV.cTnT. S1-TM-HA were digested with EcoRI and PstI, treated with Klenow fragment, and self-ligated to remove the cTnT promoter; ii) from these non-cTnT promoter plasmids, the S1-HA, S2-HA, and S1-TM fragments were cloned into pAAV.CAG vector with BamHI and SalI. The cloning primer sequences were listed in Supplemental Table 1.

#### Co-immunoprecipitation

Cell soluble protein extracts for co-immunoprecipitation were prepared in lysis buffer (20 mM Tris HCl (pH 8), 137 mM NaCl, 10% glycerol, 1% Triton X-100, 2 mM EDTA). Protease inhibitor cocktail (Roche) was added to the lysis buffer immediately before use. The protein solution was diluted with 1 volume of IP buffer (Lysis buffer without glycerol). Antibody or IgG was added to the protein extract, and antibody bound protein complexes were pulled down with pre-equilibrated protein G Dynabeads. After three washes, the immunoprecipitated proteins were eluted with 1xSDS loading buffer.

#### AAV9 packaging and administration

AAV9.cTnT.GFP and AAV9.cTnT.iCre (Lin et al., 2015) was packaged in 293T cells with AAV9:Rep-Cap and pHelper (pAd deltaF6, Penn Vector Core) and purified and concentrated by gradient centrifugation. AAV titer was determined by quantitative PCR, in which a primer pair amplifying a fragment of the chicken cardiac TnT (cTnT) promoter was used. The primers sequences were : GCTTTCACATGACAGCATCTGGGG; CCCAAGCTATTGTGTGGCCT.

For retro-orbital AAV injection, mice were anesthetized with 3% isoflurane. A 30 gauge needle was inserted at a 45° angle to the eye, lateral to the medial canthus, through the conjunctival membrane. The needle was positioned behind the globe of the eye in the retrobulbar sinus. Less than 100 μl of viral solution was injected.

Echocardiography (Echo) measurements

Before being tested, the mouse chest was depilated and the mice were trained for echo for three consecutive days. Briefly, the mice were held in a position to expose the chest, and an artificial plastic probe was put onto the chest to mimic the action of the ultrasound probe. On the third day, conscious mice were subjected to Echo measurements. The ultrasound probe and gel was applied to the chest to obtain echocardiography measurements of the heart. This typically took 5-10 minutes, during which mice were held for approximately one-minute intervals. After completion of the study, the gel was wiped from the chest with a paper towel, and the mouse was returned to normal housing.

Histology and Immunohistochemistry staining

For Hemotoxylin and Eosin (H&E) staining, hearts were fixed with 4% PFA and embedded in parafin. For chromogenic immunohistochemistry staining, fixed hearts were embedded in OCT and cryoprotected with 30% sucrose. 8-μm paraffin sections were used for H&E and immunohistochemistry staining. Primary antibodies used for this study were summarized in Supplementary Table 3. Signals were detected using the Anti-Rabbit IgG (alkaline phosphatase) Polymer Detection Kit (Vector Laboratories). Imaging was performed on Keyence microscope.

**Supplemental Table 1. Cloning primers**

| Amplicon | Forward | Reverse | Restriction enzyme sites |
|---|---|---|---|
| CoV-2-S1 | aacgctagcaggccgcctgggcccgttaacaccatg | caaccatggcggagagttggtttgggtctggtaagaa | NheI, NcoI |
| CoV-2-S2 | aacgctagcaacaccatgtccgtagccagtcaaagcataattgcgtacaccatg | acgttactagtttactaagcgtaatct | NheI, SpeI |
| TM/IC region | caaccatggccctggtacatttggctgggcttcatcgct | cacGTCGACacgttactagtttactaagcgtaatct | NcoI, SalI |
| mTLR4-LRR | CGACAAGCTTGCGGCCGCGAATTCA | accGGATCCaaaatgttgcagtattcctttagat | NotI, BamHI |
| mTLR4-TIR | aatGCGGCCGCtccaaagagtctagccttcttcaatctta | GCCACCCGGGATCCTCTAGAGTCG | NotI, BamHI |
| HA tag | catggtacccatacgatgttccagattacgctg | tcgacagcgtaatctggaacatcgtatgggtac | NcoI, SalI |

**Supplementary Table 2. qRT-PCR primers**

| Species | Gene name | Forward | Reverse |
|---|---|---|---|
| Mouse | *Myh6* | CTCTGGATTGGTCTCCCAGC | GTCATTCTGTCACTCAAACTCTGG |
| Mouse | *Nppa* | CACAGATCTGATGGATTTCAAGA | CCTCATCTTCTACCGGCATC |
| Mouse | *Il-6* | CACTTCACAAGTCGGAGGCT | CTGCAAGTGCATCATCGTTGT |
| Mouse | *Il1b* | TGTGCAAGTGTCTGAAGCAGCTA | TCAAAGGTTTGGAAGCAGCCCT |
| Mouse | *Gapdh* | CAGGTTGTCTCCTGCGACTT | GGCCTCTCTTGCTCAGTGTC |
| Mouse | *Ccl2* | GTTGGCTCAGCCAGATGCA | AGCCTACTCATTGGGATCATCTTG |

**Supplementary Table 3. Antibodies**

| Antigen | Host | Vendor (Cat #) | Usage (dilution) |
|---|---|---|---|
| Mac-3 | Rat | BD (553322) | IHC (1:100) |
| Flag | Mouse | Sigma (F1804) | Western blot (1:1000) |
| SARS-CoV-2 Spike | Rabbit | Rockland (200-401-MS9-0.1) | Western blot (1:1000)<br>IHC (1:200) |
| RelA | Rabbit | Santa Cruz (sc-372) | Western blot (1:1000) |
| His Ab | Rabbit | CST (2365S) | Western blot (1:1000) |
| HA tag | Rabbit | CST (C29F4) | Western blot (1:1000) |
